# Chemogenetic activation induces persistent transcriptomic remodeling and functional plasticity in cultured astrocytes

**DOI:** 10.64898/2026.01.20.700482

**Authors:** Momono Segawa, Hajime Yamamoto, Kentaro Abe

## Abstract

Astrocytes are essential regulators of neuronal function and brain homeostasis. Recent methodological advances have increasingly revealed their active roles through precise manipulation of astrocyte function in the brain. While chemogenetic strategies, such as Designer Receptors Exclusively Activated by Designer Drugs (DREADDs), have been widely used to recapitulate endogenous G-protein-coupled receptors (GPCRs) signaling and selectively modulate astrocyte activity, the molecular mechanisms accompanying DREADD activation remain incompletely understood. Here, we characterized the transcriptional responses of cultured cortical astrocytes and found that DREADD activation induces profound changes in astrocytic gene expression profiles. Activation of Gi-, Gq-, and Gs-coupled DREADDs elicited distinct transcriptomic responses. Transcription factor activity profiling further revealed that each DREADD subtype selectively modulates a distinct set of transcription factors that collectively shape astrocytic transcriptomic responses, thereby indicating subtype-specific transcriptional mechanisms. Importantly, DREADD activation induced persistent changes in the astrocytic transcriptome that remained detectable even three days after stimulation. These transcriptomic alterations were associated with sustained changes in astrocytic calcium responses to extracellular ATP, indicating plastic changes in astrocytic physiological function. Together, these findings provide a molecular framework for interpreting astrocytic DREADD manipulations and reveal a mechanistic basis for functional plasticity and heterogeneity of astrocytes.

## Introduction

Astrocytes are a major glial cell type and constitute one of the predominant cellular populations in the brain. Once regarded as passive or silent cells, recent technological advances have uncovered the rapid and dynamic nature of both intracellular and intercellular signaling in astrocytes, highlighting their active roles in the regulation of diverse brain functions (1–5). Experimental approaches that manipulate these dynamic signaling pathway have been employed to investigate the causal roles of astrocytic signaling molecules in neural information processing (6–8). Among the diverse signaling receptors expressed in astrocytes, G-protein-coupled receptors (GPCRs) have gathered particular attention. GPCRs are abundantly expressed in astrocytes, with hundreds of distinct subtypes expressed in an astrocyte identity- and brain region-dependent manner (9–12). Experimental manipulation of astrocytic GPCR signaling has been enabled through the use of Designer Receptors Exclusively Activated by Designer Drugs (DREADDs) (13), a chemogenetic approach that allows reversible and repeatable control of specific signaling pathways in defined cell populations with minimal invasiveness (14). DREADDs technology utilizes mutated variants of endogenous GPCRs, specifically the muscarinic acetylcholine receptors, to selectively and artificially activate intracellular G-protein signaling pathways (13). Similar to their application in neurons, DREADDs have been widely adopted in astrocyte research, where they have been instrumental in revealing essential roles of astrocytic GPCR signaling in learning, arousal, instinctive behaviors, as well as disease pathology (15–17).

Unlike in neurons, where DREADD activation typically leads to enhancement or suppression of action potential firing, the downstream consequences of DREADD activation in astrocytes are less understood and appear to be more complex (17). It has been reported that both Gi-coupled and Gq-coupled receptors activate calcium signaling in astrocytes, but in different extents and through distinct mechanisms (18, 19). Beyond calcium, DREADD activation can also elicit changes in other second messengers, such as cAMP, DAG, and IP_3_, and modulate the activity of various kinases, depending on the specific receptor activated (20–22). Importantly, these effects are often context- and brain region-dependent, reflecting the local cellular environment and signaling repertoire of astrocytes. This complex, pathway- and context-dependent signaling landscape complicates the development of a unified understanding of the molecular consequences of DREADD activation in astrocytes. An additional and often overlooked aspect of DREADD activation is its long-term impact. To date, most studies have focused on immediate effects of DREADD receptor activation, which occurs in timescales ranging from seconds to minutes following the ligand stimulation. However, GPCR activation potentially induces long-lasting changes in cell physiology (23, 24), some of which may involve persistent alterations in gene expression profiles (25). Indeed, long-term changes in gene expression have been observed in astrocytes during memory formation, a phenomenon referred to as astrocytic plasticity or astrocyte engrams (2, 25, 26).

To address this gap and better understand plastic changes in astrocytes elicited by GPCR signaling, we investigated transcriptomic alterations induced by DREADD activation and the molecular mechanisms underlying these changes. Using purified populations of cortical astrocytes in vitro, our analysis focused on the transcriptomic consequences of activating three widely used DREADD subtypes: the Gi-coupled hM4Di, the Gq-coupled hM3Dq (13), and Gs-coupled rM3Ds (27). We employed transcription factor (TF) activity reporters (28) to directly assess the impact of DREADD activation on transcriptional regulation. By integrating transcriptomic profiling with TF activity measurements, we sought to delineate the molecular programs engaged by DREADD activation and to elucidate the mechanisms underlying long-lasting changes in astrocytic state following GPCR activation.

## Results

### Purified cultured astrocytes and expression of DREADD receptors

To assess transcriptomic changes induced by DREADD activation in astrocytes, we prepared astrocyte monocultures from the cortices of neonatal mice (*Mus musculus*) following established protocols (29, 30). Immunostaining of these cultures revealed high astrocytic purity compared with primary cortical cultures, which contain neurons, astrocytes and other glial cell types (Fig. 1A). The majority of cells expressed astrocytic markers, such as GFAP (86.6%) and S100B (81.8%), whereas markers for neurons (NeuN/FOX3, 0.0%), oligodendrocytes (MBP, 6.9%), and microglia (IBA1, 0.007%) were negligible (Fig. 1B). Quantitative RT-PCR analysis further confirmed that, in addition to GFAP and S100B, the cultures expressed astrocyte-enriched genes including *Serpine2*, *Sparc*, *Apoe*, *Sphk1*, *Lrp1*, and *Timp3*, whereas the microglial marker *Cx3cr1* was not detected (Fig. 1C). Astrocytes are known to exhibit differential gene expression patterns across the brain regions (31, 32). In our astrocyte cultures, we observed enrichment of genes previously associated with cortical astrocytes, such as *Sparc*, *Prelp, Sphk1*, *Ccnd1*, and *Gnas* (31), relative to primary cortical cultures (Fig. 1C). Collectively, these results indicate that the astrocyte monocultures exhibit molecular characteristics of mature cortical astrocytes. In this culture, we expressed three subtypes of DREADD receptors—hM4Di, hM3Dq and rM3Ds—by plasmid transfection and stimulated them with deschloroclozapine (DCZ), which does not require metabolic conversion to exert its effect (33) (Fig. 1D).

**Fig. 1.**
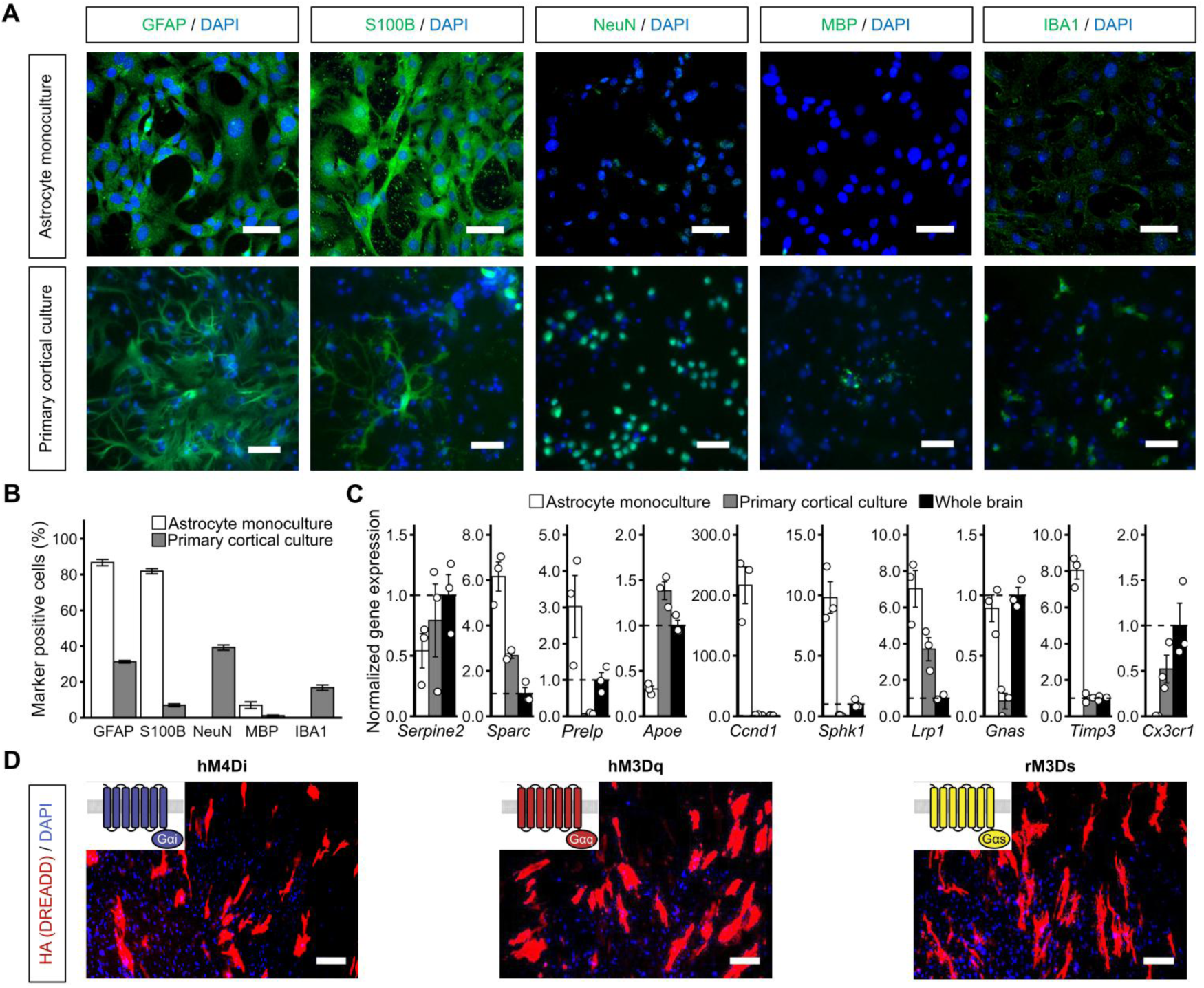
Molecular evaluation of astrocyte monoculture. (**A**) Images of immunofluorescence staining of astrocyte monoculture and primary cortical culture. Scale bar, 50 μm. (**B**) Ratio of GFAP, S100B, NeuN, MBP, and IBA1 positive cells against DAPI positive nucleus. Data are presented as mean ± SEM. Sample numbers: astrocyte monoculture, 6–9 wells from 3 independent experiments; primary cortical cultures: 4–9 wells from 2 independent experiments. (**C**) Quantitative RT-PCR analysis of astrocyte-enriched genes. Expression levels were normalized with those of GAPDH and scaled to the mean expression level of RNAs from whole mouse brain for each gene. Data are presented as mean ± SEM. Sample numbers: astrocyte monoculture, 3; primary cortical culture, 3; whole brain, 3. (**D**) Immunofluorescence staining of astrocyte monocultures expressing hM4Di, hM3Dq, and rM3Ds. Receptor expressions are visualized with HA-tag on their N-terminus. Scale bar, 100 μm.

### DREADD activation induces subtype-specific alterations in astrocyte transcriptome

We first examined transcriptomic changes induced by astrocytic DREADD activation, extending beyond previously characterized immediate signaling responses. To define transcriptional signatures associated with DREADD activation, we stimulated cultured astrocytes expressing hM4Di, hM3Dq, or rM3Ds with DCZ (11 μM) and collected RNA samples 90 min after stimulation for RNA-seq analysis (Fig. 2A). To assess global differences in the transcriptomic landscape, we defined differential expressed gene (DEG) relative to EGFP-transfected controls using criteria of *p* < 0.05 and an absolute log_2_ fold change > 1. Under these criteria, we identified 153 DEGs for hM4Di, 365 for hM3Dq, and 173 for rM3Ds (Fig. 2B). Transcriptomic analysis further revealed distinct gene expression profiles for each of the three receptor subtypes (Fig 2C). Notably, the DEG sets showed minimal overlap among the three receptor subtypes. Among the 593 genes classified as DEGs in at least one subtype, overlap among subtypes was limited. Specifically, 10.3% were shared between hM4Di and hM3Dq (61 genes; upregulated, 2.4%; downregulated, 7.9%), 2.5% between hM4Di and rM3Ds (15 genes; upregulated, 1.3%; downregulated, 1.2%), and 5.1% between hM3Dq and rM3Ds (30 genes; upregulated, 3.7%; downregulated, 0.7%; oppositely regulated between subtypes, 0.7%). Only 8 genes (upregulated, 0.67%; downregulated 0.67%) overlapped across all three DREADD subtypes (Fig. 2D).

**Fig. 2.**
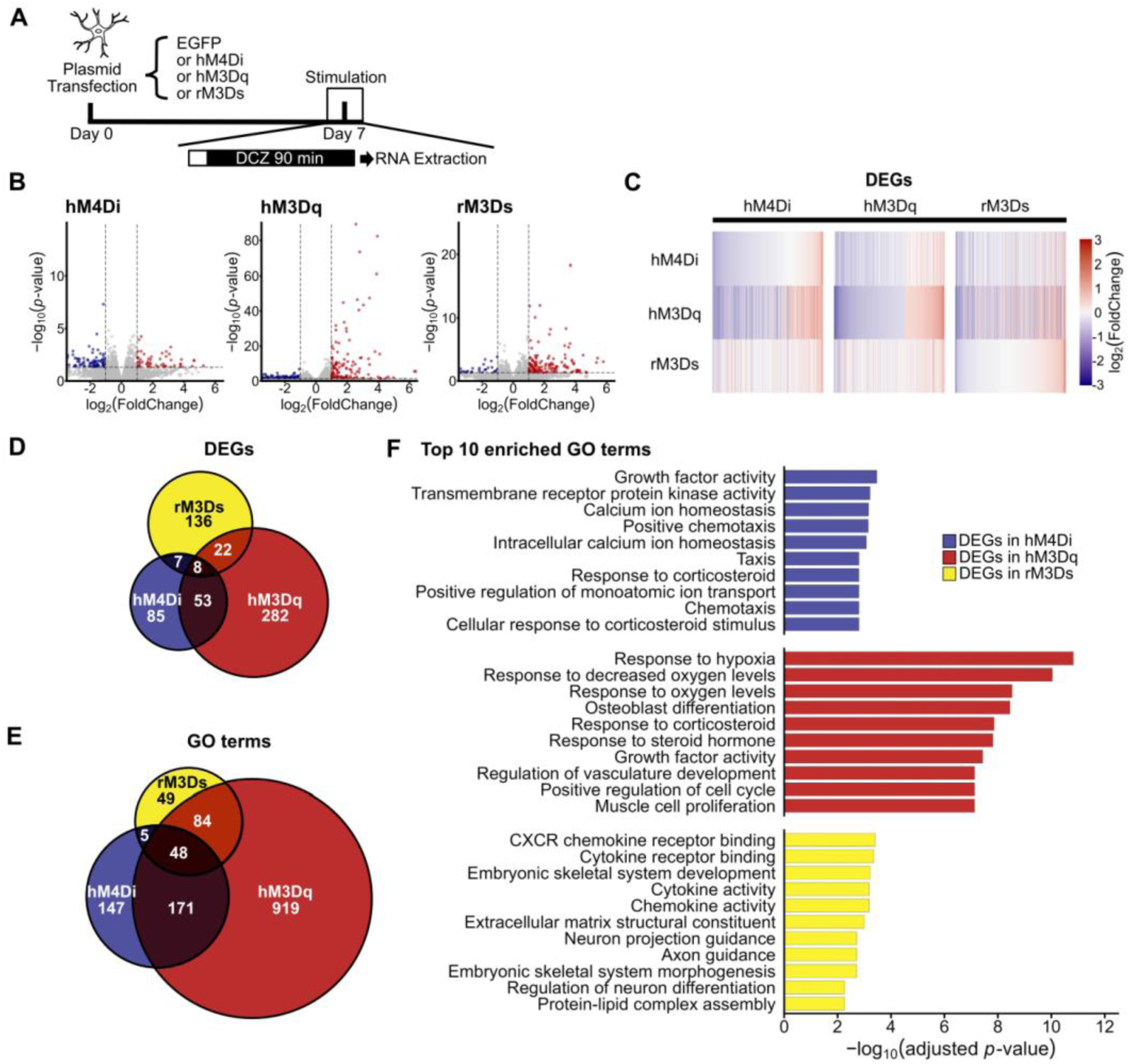
DREADD activation elicits transcriptomic changes in astrocytes. (**A**) Schematic illustration of the experimental timeline. (**B**) Volcano plots showing gene expression changes 90 min after DCZ stimulation in astrocytes expressing each DREADD subtype. (**C**) Heatmaps illustrating expression changes of DEGs identified for each DREADD subtype. Columns show DEGs defined for hM4Di, hM3Dq, and rM3Ds (criteria: *p* < 0.05 and an absolute of log_2_ fold change > 1), Rows show log_2_ fold changes of these genes following activation of the indicated DREADD subtype. (**D**) Euler diagram displaying the relationships among DEGs defined by activation of hM4Di (blue), hM3Dq (red), and rM3Ds (yellow). (**E**) Relationship among enriched GO terms for hM4Di, hM3Dq, and rM3Ds. (**F**) Top ten enriched GO terms with the lowest adjusted *p*-value associated with DEGs from each DREADD- activated culture. The top ten GO terms with the lowest adjusted *p*-value in the “Biological process” and “Molecular function” categories are shown.

Next, to further elucidate the biological significance of the transcriptomic changes observed across the three DREADD subtypes, we performed Gene Ontology (GO) enrichment analysis of the DEGs identified for each subtype. This analysis revealed subtype-specific GO terms that were significantly enriched (adjusted *p* < 0.05) across the three DREADD subtypes (Fig. 2E). Among the 1,423 GO terms significantly enriched in the DEGs of at least one subtype, 10.3% were unique to hM4Di, 64.6% to hM3Dq, and 3.4% to rM3Ds. Only 48 GO terms were commonly enriched across all three DREADD subtypes, representing 12.9% of the GO terms enriched in hM4Di, 3.9% in hM3Dq, and 25.8% in rM3Ds (Fig. 2E). Specifically, DEGs associated with hM4Di activation were enriched for terms related to calcium ion homeostasis and signal transduction, whereas those associated with hM3Dq activation were enriched for responses to hypoxia, extracellular stress, and cell proliferation, and those associated with rM3Ds activation were enriched for extracellular matrix, axon guidance, and chemokine/cytokine activity (Fig. 2F). Taken together, these results indicate that DREADD stimulation induces substantial transcriptional responses in astrocytes. The largely distinct sets of genes altered across the three receptor subtypes points to Gi-, Gq-, and Gs-specific signaling, rather than a nonspecific response to ligand application. Furthermore, GO enrichment analysis suggests that these transcriptional changes may underlie subtype-specific alterations in astrocytic physiology.

### Differential activation of transcription factors induced by DREADD activation in cultured astrocytes

TFs directly regulate the transcription of their target genes. To gain mechanistic insight into the differential gene expression induced by the three DREADD subtypes, we sought to identify the TFs that may mediate the transcriptional regulation of their respective DEGs. To this end, we first performed promoter analysis of DEGs to uncover candidate TFs potentially involved in their regulation. Utilizing RNA-seq data, we examined enrichment of transcription factor binding sequences (TFBSs) within promoter regions (±1,000 bp from transcription start sites) of DEGs for each DREADD subtype (Fig. 3A). This analysis revealed largely distinct sets of TFBSs specifically enriched in each DREADD subtype. These findings suggest that subtype-specific TF activity downstream of DREADD activation underlies the observed divergence in transcriptional regulation.

**Fig. 3.**
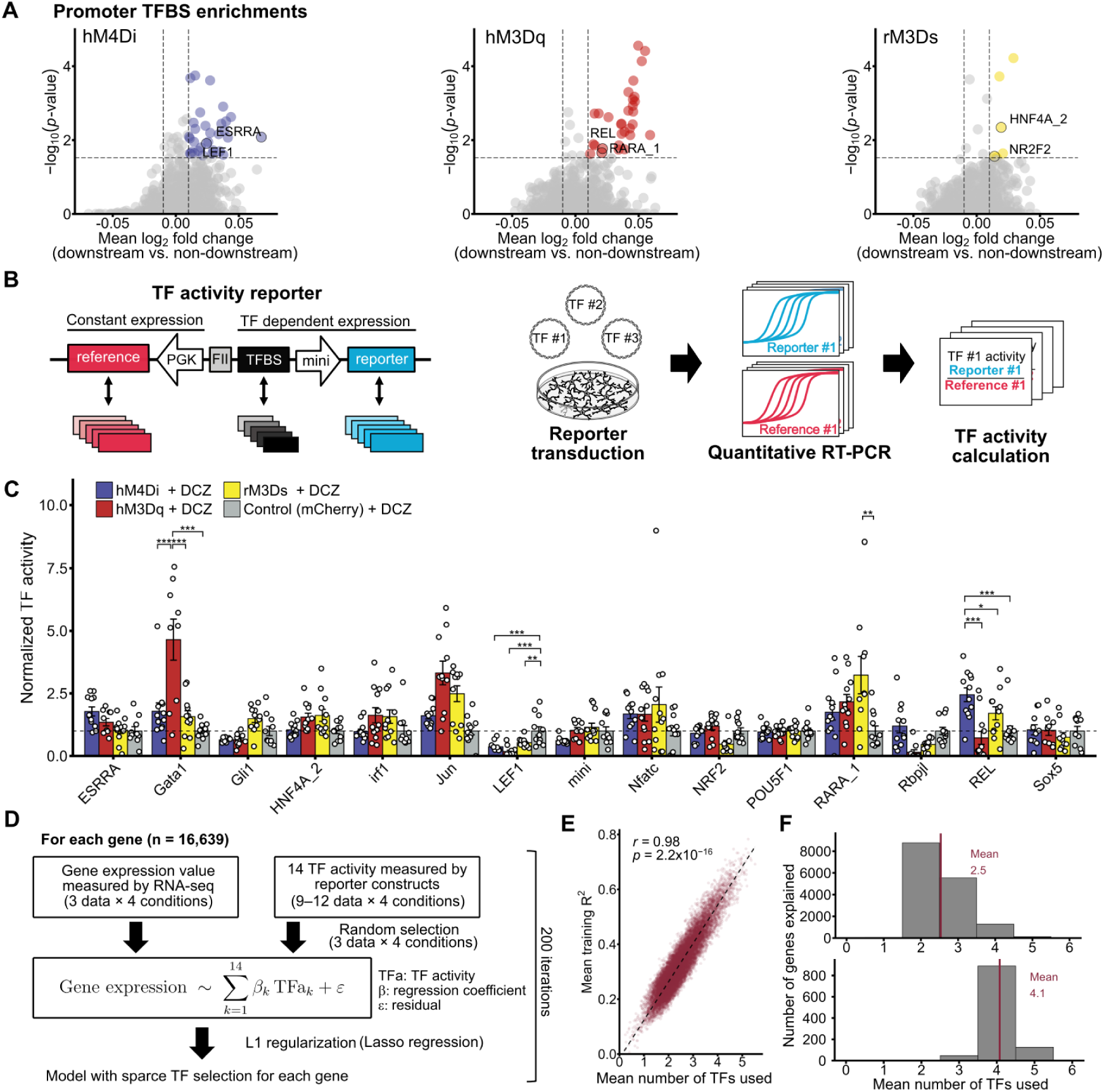
DREADD activation results in distinct TF-activity patterns. (**A**) Mean log_2_ fold change of TFBS enrichment in downstream versus non-downstream genes and the corresponding statistical significance (−log_10_ *p*-value). Dotted lines indicate thresholds (log_2_ fold change = ±0.01, and *p* = 0.03). (**B**) Schematics of TF activity profiling. (**C**) Normalized TF activity across each DREADD subtype and control conditions. Mean ± SEM; one-way ANOVA, Tukey’s post hoc test; *n* = 9–12 samples per group. *p* < 0.05 (*), *p* < 0.01 (**), and *p* < 0.001 (*), adjusted for multiple comparisons. (**D**) Overview of the multiple linear regression framework. (**E**) Relationship between the mean training *R*^2^ and the mean number of TFs used in the model. Each dot represents a single gene (total *n* = 16,639). Pearson correlation coefficient is shown. (**F**) Distribution of genes whose expression levels are explained by a given number of TFs analyzed in the reporter assay. Red lines indicate the mean number of TFs used across all explained genes. For each gene, the mean number of TFs used across 200 iterations was rounded to the nearest integer.

Next, to assess whether DREADD activation affects the gene transcriptional activity of TFs, we directly measured the activity of candidate TFs. Specifically, cultured astrocytes were co-transfected with a DREADD expression plasmid (either hM4Di, hM3Dq, rM3Ds, or mCherry as a control) together with 3–6 distinct TF activity reporter plasmids, each targeting a different TF. These reporter plasmids contain TFBSs upstream of a synthetic minimal promoter, enabling quantification of TF activity by comparing reporter gene expression under control and stimulated conditions (Fig. 3B). Among the candidate TFs implicated by promoter analysis of DEGs (Fig. 3A), we measured the activity of 15 TFs using previously established TF activity reporter constructs (28). This analysis revealed subtype-specific changes in TF activity following activation of each of the three DREADD subtypes. Specifically, GATA activity was enhanced upon hM3Dq activation among other DREADD subtypes, REL for hM4Di and RARα for rM3Ds specific activation (Fig. 3C). We also observed reduced activity of TCF/LEF across all three DREADD subtypes compared with control culture (Fig. 3C). These findings indicate that DREADD activation alters TF activity, which is associated with subsequent changes in gene expression. Moreover, distinct sets of TFs are differentially modulated across three DREADD subtypes.

Do differential changes in TF activities account for the distinct gene expression patterns observed across DREADD subtypes? To address this question, we adopted a theoretical modeling approach. We constructed linear regression models to explain the expression level of each gene using multiple TF activities, with the number of TFs constrained by LASSO regularization. Specifically, the models utilized TPM value for each of the 16,639 genes quantified by RNA-seq and the activity of 14 TFs (excluding minimal promoter) measured in our reporter assays across the activation of three DREADD subtypes and control condition (Fig. 3D). As expected, we observed a significant positive correlation between the training *R*^2^ and the number of TFs included in the model (Fig. 3E; Pearson’s correlation = 0.98, *p* < 2.2 × 10^−16^), indicating that the models using combinatory TF activities more effectively capture gene expression landscapes elicited by DREADD activation. For example, expression levels of 94.9% of genes were explained at a threshold of *R*^2^ > 0.2, with a mean of 2.9 TFs used per model. A higher threshold of *R*^2^ > 0.5 was achieved for 6.3% of genes, requiring a mean of 4.1 TFs per model (Fig. 3F). This observation—incorporating multiple TFs increases the explanatory power of the model—is consistent with the fact that promoters in human genome contain an average of 5.7 TF binding sites (34), indicating that multiple TFs act synergistically in the dynamic regulation of individual gene expression.

Together, these results indicate that activation of each DREADD subtype induces a distinct pattern of TF activity, which in turn contributes to the emergence of subtype-specific transcriptomic changes.

### DREADD activation induces plastic changes in astrocyte transcriptome

Given that DREADD activation drives distinct changes in TF activity (Fig. 3B), we next asked whether these signaling events induce long-lasting alterations in the astrocytic transcriptome, thereby shifting astrocytic state. To address this, we tested whether the transcriptomic alteration triggered by DREADD activation remained detectable after an extended period following ligand removal. Specifically, cultured astrocytes expressing hM3Dq were stimulated with DCZ as described above, after which the medium was replaced with ligand-free medium. We then performed RNA-seq analysis on 3 days after the stimulation and compared these results with those obtained 90 min after stimulation (Fig. 4A). To this experiment, we adopted more stringent definition of DEGs using the criteria of an adjusted *p* < 0.05 and an absolute log_2_ fold change > 1. Under these criteria, we identified 36 DEGs in astrocytes collected 3 days post-stimulation, while identifying 86 DEGs in those collected 90 min after stimulation (Fig. 4B). Comparison of the two time points revealed both differences and commonalities in gene expression changes (Fig. 4C). We observed that DEGs detected at the two post-DCZ time points were largely distinct, with only 2 genes (1.7%) shared between the two DEG sets (Fig. 4D). Gene Ontology (GO) enrichment analysis of these DEGs revealed that, at the earlier timepoint, terms related to hormonal responses and cell proliferations were enriched. In contrast, at the later timepoint, GO terms related to functional changes of the brain exhibited enrichment, such as “Adult behavior” or “Neurotransmitter uptake”, suggesting that Gq-DREADD activation induces shifts in the physiological properties of astrocytes at later stages (Fig. 4E). We next assessed shared gene expression changes between the early and late timepoints by utilizing Rank-Rank Hypergeometric Overlap (RRHO) analysis, which provides a threshold-free evaluation of gene sets commonly regulated across two experimental conditions (35). Unlike DEG analysis, this threshold-free analysis identified 1,462 commonly upregulated genes and 1,276 commonly downregulated genes between the two time points (Fig. 4F). Gene Ontology (GO) analysis of the commonly upregulated gene sets revealed significant enrichment in terms related to protein translation such as “cytoplasmic translation”, “translation at synapse”, and “cellular respiration”. In contrast, the commonly downregulated gene sets were significantly enriched in terms related to cell division such as “chromosome segregation”, “sister chromatid segregation”, and “nuclear chromosome segregation” (Fig. 4G). Taken together, these results indicate that DREADD activation evokes astrocytic gene expression changes that remain detectable even three days after stimulation, suggesting sustained transcriptional reprogramming.

**Fig. 4.**
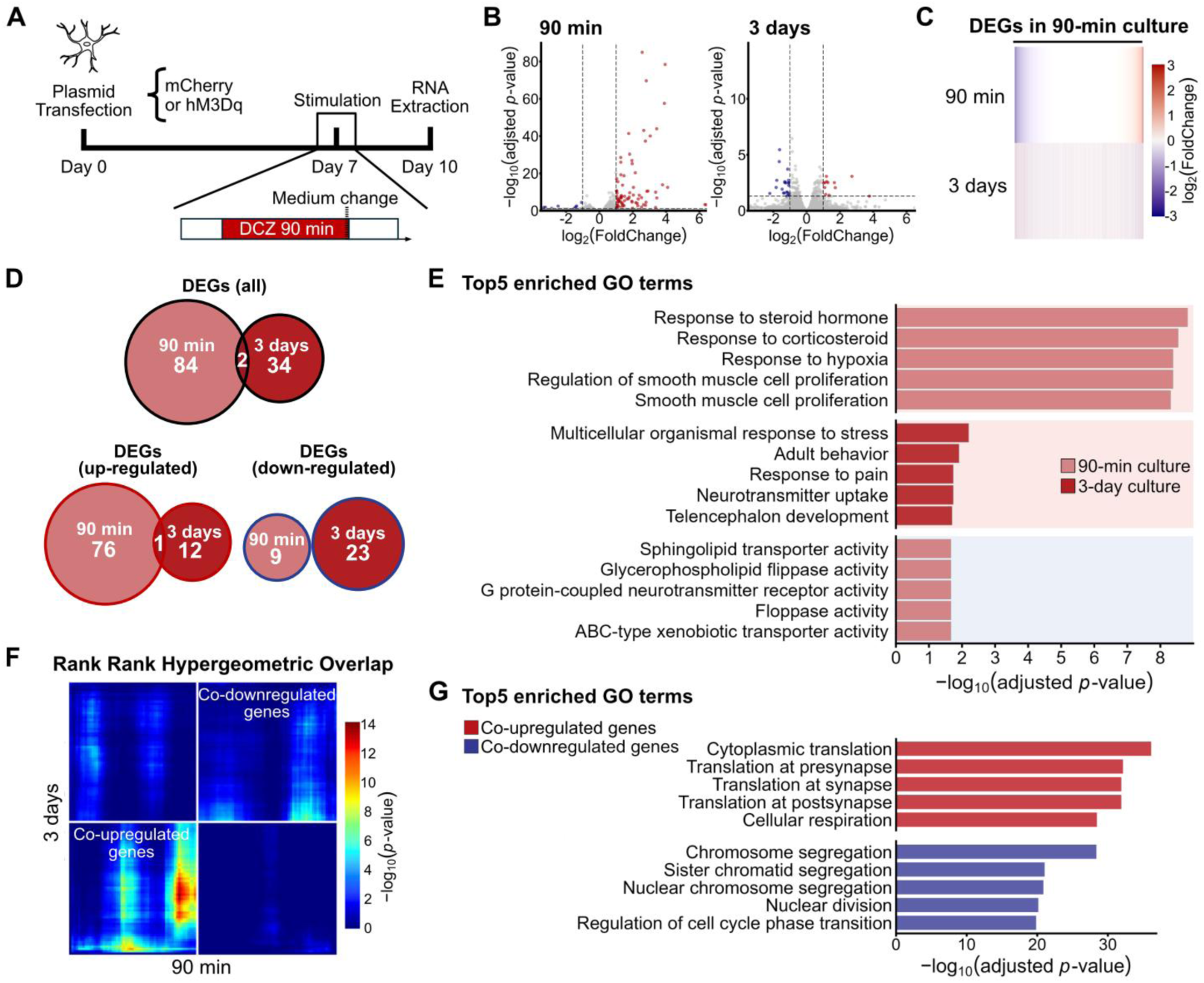
Plastic gene expression changes induced by Gq-DREADD activation. (**A**) Schematic illustration of the experimental timeline. (**B**) Gene expression changes at 90 min and 3 days after DCZ stimulation in astrocytes expressing Gq-DREADD. (**C**) Gene expression changes observed in 90 min and 3 days after DCZ stimulation. Only genes identified as DEGs for 90 min conditions are shown. (**D**) Relationship between DEGs identified for 90-min and 3-day conditions. All DEGs, up-regulated DEGs (red), and downregulated DEGs (blue) are shown separately. (**E**) GO enrichment analysis for significantly up- or down-regulated genes at 90-min and 3-day after activation of Gq-DREADD. The top five GO terms with the lowest adjusted *p*-values in the “Biological process” and “Molecular function” categories are shown. Red and blue backgrounds denote GO terms enriched in upregulated and downregulated DEGs, respectively. (**F**) RRHO2 map comparing gene expression changes in astrocytes between 90-min and 3-day post-stimulus conditions. Pixels in the heatmap represent the significance of overlap. (**G**) GO analysis of genes commonly upregulated or downregulated as defined by RRHO analysis. Top five GO terms with the lowest adjusted *p*-value in the “Biological process” are shown.

### DREADD activation induces long-lasting functional changes

It is well established that astrocytes respond to extracellular ATP (36). In our transcriptomic analysis on astrocytes 3 days after a 90-min DCZ stimulation, we noticed a collective downregulation of ATP receptor genes detected in RNA-seq (*P2rx2*, *P2rx3*, *P2rx4*, *P2rx5*, *P2rx6*, *P2rx7*, *P2ry1*, *P2ry2*, *P2ry4*, and *P2ry6*) (Fig. 5A, B; *p* = 0.013, LME: fixed effect, condition; random effect, sample and gene). Therefore, we next assessed whether this transcriptional downregulation resulted in a functional alteration of the cellular responsiveness to ATP. Using the genetically encoded calcium indicator GCaMP6s, which was transfected together with hM3Dq, we investigated changes in intracellular calcium concentration in response to extracellular ATP (Fig. 5C). In unstimulated control cultures, ATP (100 μM) elicited a robust, transient increase in cytosolic Ca^2+^ (Fig. 5D). In contrast, 3 days after a brief 90-min DCZ stimulation, astrocytes exhibited a significant reduction of responsiveness to extracellular ATP (Fig. 5D). Both the percentage of ATP-responsive cells (Fisher’s exact test, *p* < 0.0001; Fig. 5E) and the integral of Ca^2+^ responses (AUC) (unpaired *t*-test, *p* = 0.004; Fig. 5F) were significantly reduced compared to unstimulated controls, whereas no detectable changes in astrocyte size were observed 3 days following DCZ stimulation. These results indicate that a single, transient DREADD activation event results in sustained functional remodeling of cultured astrocytes.

**Fig. 5.**
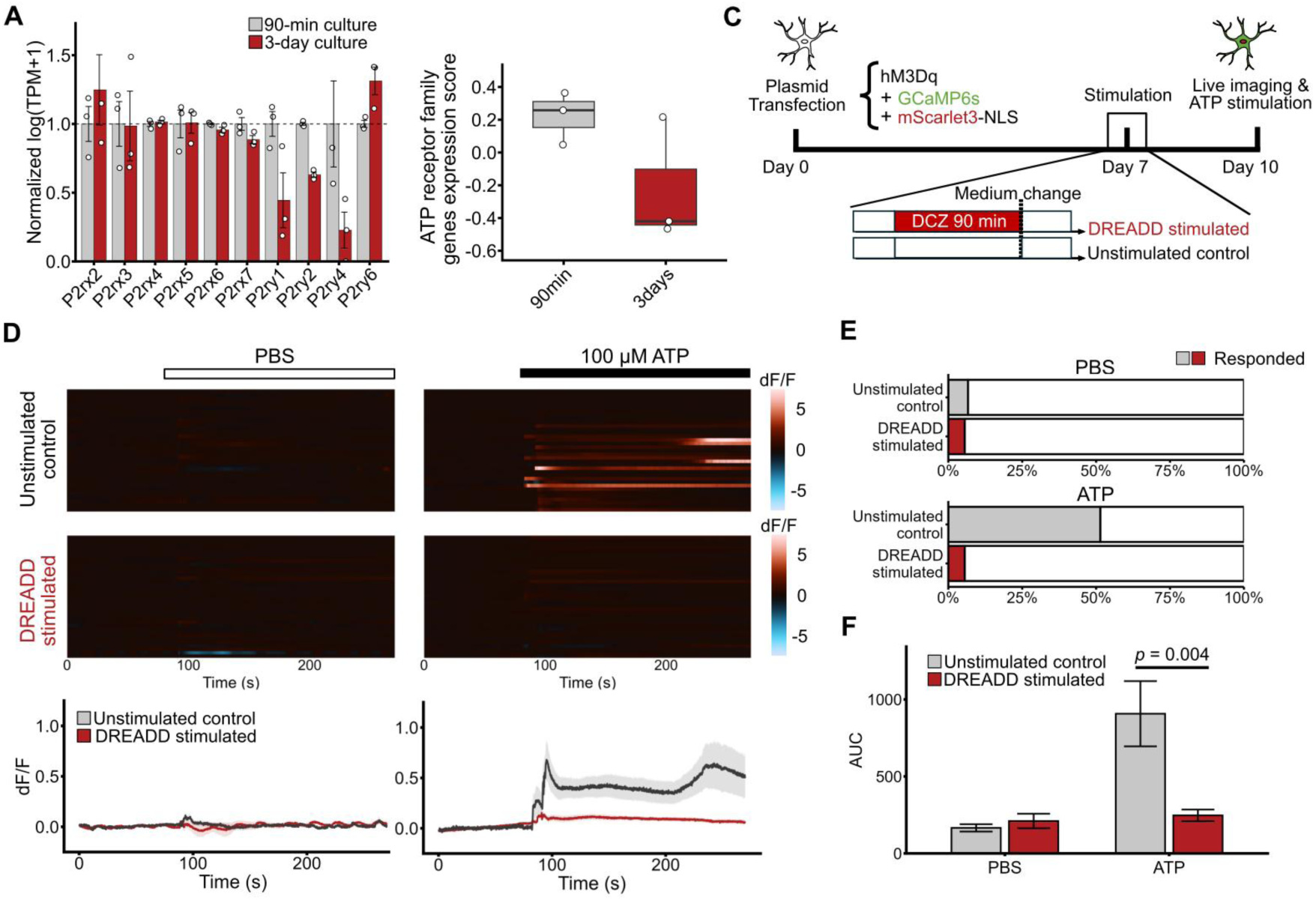
Gene expression changes induced by Gq-DREADD activation leads to sustained functional alterations in astrocytes. (**A**) Expression changes of ATP receptor family genes in cultured astrocytes measured immediately after (gray) or 3 days after (red) 90-min hM3Dq activation. Relative expression levels normalized to the 90-min culture are shown. Data are presented as mean ± SEM, *n* = 3 samples per group. All detected ATP receptor family genes (P2X and P2Y receptors) identified by RNA-seq are shown. (**B**) Z-scored log (TPM + 1) for ten ATP receptor family genes shown in (**A**) are averaged for each sample. (**C**) Schematic illustration of calcium imaging experiments assessing Ca^2+^ responses to ATP stimulation. (**D**) Heatmaps (top) and ΔF/F traces (bottom) of astrocytic Ca^2+^ responses. Solid lines and shaded areas indicate mean ± SEM. ATP (100 μM) or PBS was applied at 90 s. (**E**) Percentage of cells exhibiting Ca^2+^ increases in response to ATP stimulation. (**F**) Average area under curve (AUC) of astrocytic calcium signals. Data are shown as mean ± SEM, *n* = 30–36 cells per group from 3 independent experiments.

## Discussion

This study revealed that DREADD activation in astrocytes elicited profound and subtype-specific alterations in gene expression profiles. We further demonstrated that the three DREADD subtypes, hM4Di, hM3Dq, and rM3Ds, each induced a distinct transcriptional response. To gain mechanistic insight into this divergence, we showed that each subtype drives a distinct pattern of TF activation, which likely underlies the observed transcriptomic differences among subtypes. The transcriptomic changes induced by Gq-DREADD activation persisted for at least 3 days following stimulation, and astrocytic responsiveness to extracellular ATP exhibited a sustained change, which correlated with the downregulation of its receptors.

Given the substantial heterogeneity of astrocytes in the brain, we employed a purified population of astrocytes in an in vitro culture platform. By focusing on a homogeneous astrocyte population, we were able to delineate downstream molecular differences elicited solely by activation of distinct DREADD subtypes. Although our analysis was conducted in vitro, the distinct gene expression responses and TF activation patterns suggest that similar subtype-specific effects may also occur in astrocytes in vivo. Astrocytes express hundreds of GPCRs, including receptors for glutamate, serotonin, adenosine, purines, and oxytocin, which couple to diverse G-proteins to mediate astrocytic responses to neuronal activity and neuromodulators (10). DREADDs are designed to artificially elicit intracellular signaling of these endogenous GPCRs, primarily to regulate neuronal activity (14). Notably, when expressed in astrocytes, it has been observed that the downstream effects of GPCRs or DREADD activation on brain functions, such as memory formation, often vary across receptor subtypes (15, 37–41). While previous work has attributed some of this variability to differences in calcium transient dynamics (17), our data extend this view by demonstrating that DREADD activation elicits divergent intracellular signaling programs that result in unique gene expression signatures. Together with previous observations, our results highlight the profound impact of GPCR subtype-specific signaling pathways on astrocyte physiology. Remarkably, we found that the gene expression changes induced by acute Gq-DREADD activation remained detectable for at least 3 days following the end of stimulation period. Analysis of DEGs revealed that the transcriptional profiles at early and late time points were largely distinct, suggesting a temporal shift in the gene sets robustly modulated by DREADD activation (Fig. 4C and D). These results suggest that DREADD activation appears to exert long-lasting effects by reorganizing gene regulatory programs rather than by simply maintaining the initial expression patterns of DEGs. These findings support the notion that DREADD activation can induce durable, pathway-specific plasticity in astrocytic states, often referred to as “astrocytic plasticity” (2, 25, 26). This concept is further supported at a functional level by our observation that astrocytic responsiveness to extracellular ATP underwent sustained modulation following Gq-DREADD activation, correlating with the downregulation of ATP receptor genes (Fig. 5). These findings align well with previous reports showing that noradrenaline-mediated GPCR stimulation in astrocytes modifies surface receptor expression, thereby immediately altering responsiveness to neurotransmitters to which cells were previously insensitive (42). In addition to this immediate effect, we observed that the phenotypic changes induced by GPCR stimulation persisted and remained evident even 3 days after stimulation. Collectively, these observations suggest that astrocytes exhibit dynamic transcriptomic flexibility by integrating extracellular signals through a vast array of GPCRs. This mechanism should allow for changes in cellular responsiveness, ensuring precise adaptation to the physiological contexts. Such persistent molecular and functional alterations in astrocytic state may also help explain the often inconsistent findings across laboratories regarding how DREADD activation influences astrocytic calcium signaling and synaptic plasticity (19).

To explore the upstream regulators of the transcriptomic changes induced by DREADD stimulation, we assessed the activity of 15 TFs selected based on binding-site enrichment in the promoters of identified DEGs. Among these, we identified specific TFs whose activity was selectively modulated by individual DREADD subtypes (Fig. 3C). It should be noted that the selection of 15 TFs was not exhaustive, and other TFs are likely to contribute. Nevertheless, the distinct TF activity patterns identified in this study provide specific molecular candidates that may underlie the observed transcriptomic divergence, as supported by our modeling approach (Figs. 3D–F). Although the specific roles of these TFs in astrocytes have not yet been fully characterized, the dysregulation of the TFs highlighted in this study—including GATA, REL, RARα, and TCF/LEF—has been linked to various brain pathologies (43–46). In addition, dysregulation of astrocytic GPCR signaling has been implicated in several neurological disorders, including Alzheimer’s disease (47), Parkinson’s disease (48), and epilepsy (49). Therefore, investigating the downstream molecular mechanisms triggered by DREADD activation can contribute not only to understanding of the functional consequences of endogenous GPCR activation in astrocytes, but also to the development of therapeutic strategies targeting astrocytic GPCR pathways in the treatment of brain disorders.

This study has several limitations. First, the use of cultured and purified astrocyte populations may not fully capture the complex and heterogeneous nature of astrocytes in the intact brain. Whether the differential molecular consequences of DREADD activation observed in this study are conserved in vivo remains an open question. Furthermore, whether these responses vary across astrocyte identity needs investigation. Second, while we demonstrate persistent transcriptional and functional changes following astrocytic DREADD activation, the long-term functional consequences of this manipulation, as well as the extent to which it recapitulates physiological GPCR signaling in the brain, remain to be fully elucidated. Addressing these issues will require detailed, multifaceted investigations spanning molecular, physiological, and behavioral levels.

Overall, this study provides a foundation for decoding the molecular basis of chemogenetic manipulation in astrocytes. We further propose that astrocytic GPCR activation induces persistent, pathway-specific changes in cellular states—a form of molecular plasticity that likely reflects their capacity to adapt to the local physiological environment.

## Materials and Methods

### Animals

The care and experimental manipulation of animals used in this study were reviewed and approved by the Institutional Animal Care and Use Committee of Tohoku University. All experiments and maintenance were performed following relevant guidelines and regulations.

### Cell cultures and transfection

Cortical astrocyte monoculture was prepared from mice (*Mus musculus*, Slc:ICR, SLC Japan) as previously described (29). Briefly, cortices were dissected from the brains of P1–P4 neonatal mice. Tissues from multiple pups of both sexes were pooled and processed together. Dissected brain tissues were treated with trypsin, and dissociated cells were plated onto cell culture dishes or 24-well plates pretreated with poly-L-lysine (Sigma-Aldrich, #P2336) at 37°C under 5% CO_2_, using high-glucose Dulbecco’s Modified Eagle Medium (DMEM, Fujifilm-Wako) supplemented with 10% fetal bovine serum (Thermo-Fisher) and penicillin-streptomycin-amphotericin B (100×; Fujifilm-Wako). Plasmid transfection was performed using HilyMax reagent (Dojindo Molecular Technologies) according to the manufacturer’s instructions. The medium was replaced to a fresh medium on the next day of transfection. Cells cultured in 10 cm dishes were used for RNA-seq experiments, while those seeded in 24-well plates were used for transcription factor activity assays, Ca^2+^ imaging, and immunostaining. For in vitro drug treatment, cultured astrocytes were treated with DCZ (deschloroclozapine, MedChemExpress, #HY-42110, used at 11 μM; or deschloroclozapine dihydrochloride, MedChemExpress, #HY-42110A, used at 8 μM) for 90 min, 7 days after plasmid transfection. Primary cortical cultures were prepared as described before (50). Briefly, neocortex was collected from mouse embryos on embryonic day 15, treated with trypsin, and plated on 24-well plates pretreated with poly-L-lysine. The cultured cortical neurons are maintained with Neurobasal plus medium (Thermo-Fisher, #A3582901) supplemented with B27 plus supplement (20×; Thermo-Fisher, #A3582801), and penicillin-streptomycin (100×; Fujifilm-Wako).

### Plasmid constructs

Plasmid used for DREADD receptor expression are following: pAAV-CMV-EGFP, pAAV-GFAP-CRY, pAAV-CMV-HA-hM3Dq, pAAV-CMV-HA-hM3Dq-IRES-SCR3-WpA, pAAV-CMV-HA-hM4Di, pAAV-CMV-HA-hM4Di-IRES-SCR3-WpA, pAAV-CMV-HA-rM3Ds and pAAV-CMV-HA-rM3Ds-IRES-SCR3-WpA. Plasmid construction was performed by conventional methods based on restriction enzyme and gene fragment synthesis: pAAV-CMV-EGFP was created from pAAV-pgk-DIO-rvCRY-WPRE (RRID: Addgene_182060) by replacing pgk-DIO-rvCRY with synthesized gene fragment of CMV-EGFP. pAAV-CMV-hM3Dq-mCherry was created from pAAV-hSyn-HA-hM3D(Gq)-IRES-mCitrine, (RRID: Addgene_58536; a gift from Bryan Roth) by subcloning HA-hM3Dq into EGFP locus of pAAV-CMV-EGFP; pAAV-CMV-hM4Di-mCherry was created from pAAV-hSyn-DIO-HA-hM4D(Gi)-IRES-mCitrine (RRID: Addgene_50455; a gift from Bryan Roth), by subcloning HA-hM4Di into EGFP locus of pAAV-CMV-EGFP; and pAAV-CMV-HA-rM3Ds was created from pAAV-GFAP-HA-rM3D(Gs)-IRES-mCitrine (RRID: Addgene_50472; a gift from Bryan Roth), by subcloning HA-rM3Ds into EGFP locus of pAAV-CMV-EGFP. Plasmid used for live imaging of intracellular calcium signals are following: pAAV-CAG-GCaMP6s, pAAV-Pgk-mScarlet3-NLSPEST, and pAAV-CMV-HA-hM3Dq. pAAV-CAG-GCaMP6s was created from pAAV-CAG-FLEX-tdTomato (RRID: Addgene_122501; a gift from Kimberly Ritola), by replacing FLEX-tdTomato by synthesized GCaMP6s with AgeI and BsrGI linker sequence; pAAV-Pgk-mScarlet3-NLSPESTwas created from synthesized gene fragment encoding human phosphoglycerate kinase 1 (Pgk) promoter, mScarlet3, and nucleus localization peptide signal (APKKKRKV) and synthetic PEST peptide sequence (NSHGFPPEVEEQAAGTLPMSCAQESGMDRHPAACASARINV). Plasmid used for TF activity measurements are referred to as LV system in a previous study (28) and following constructs were used: pLV-TFRep2-RBP, pLV-TFRep2-RAR, pLV-TFRep1-GATA, pLV-TFRep1-ESR, pLV-TFRep1-GLI1, pLV-TFRep3-TCF/LEF, pLV-TFRep4-HNF4, pLV-TFRep4-NFAT, pLV-TFRep4-STAT1/2, pLV-TFRep4-ap1, pLV-TFRep5-SOX9, pLV-TFRep5-POU5F1, pLV-TFRep5-mini, pLV-TFRep6-RELB, and pLV-TFRep6-NRF1 (28). These vectors convey two expression cassettes in single plasmid: transfection reference genes under constitutive PGK promoter, and reporter gene under transcription factor activity dependent promoter. pTFRep1 to pTFRep6 contains distinct set of reference genes and reporter genes and can be used in combination to assess up to six TF activity independently (28). All constructed plasmids were verified by Sanger sequencing.

### Immunohistochemistry

Astrocyte monocultures and primary cortical cultures (∼14 days in vitro) were fixed with 4% paraformaldehyde (Nacalai tesque) in PBS and then permeabilized with 0.25% Triton X-100 / PBS for 20 minutes, blocked with 10% Normal Goat Serum (Biowest, #S1810-500) for 20 minutes, and stained by sequential incubation with primary antibodies diluted with PBS at 4°C overnight and with Alexa Fluor 488- or Alexa Fluor 555-conjugated secondary antibodies (1:450; Thermo-Fisher) with 4’,6-diamidino-2-phenylindole (DAPI; 1:1000, Dojindo Molecular Technologies) for an hour. The primary antibodies used were as follows: anti-GFAP (1:1000; Biolegend, #PRB-571C), anti-MBP (1:500; BioLegend, #836504), anti-S100B (1:1500; Proteintech, #66616-1-Ig), anti-IBA1 (1:1000; WAKO, #019-19741), and anti-NeuN (1:1000; BioLegend, #sc374412), and anti-HA (1:1000; MBL, #M180-3MS). Fluorescent images were obtained using fluorescent microscopies with a 20× objective lens (Axiovert A1 or Axio Imager 2, Zeiss) and a 4 × objective lens (BZ-9000 or BZ-810, Keyence). The number of cells were quantified by ImageJ (NIH ImageJ) and EBImage package (v4.4.2) in R.

### Transcriptome analysis

Total RNA was collected from cultured astrocytes 90 min after DCZ application, using Phenol-chloroform based RNA extraction. For the experiments aimed at assessing the long-term effect of DREADD activation, the cells were treated with DCZ for 90 min and the medium was replaced to a fresh medium without DCZ. Then, the RNA was collected from the culture 3 days after. Purified total RNA samples underwent mRNA purification, poly-A mRNA enrichment library preparation, and sequencing on MGI DNBSEQ-T7 platform (150 bp paired-end), performed by Novogene. The raw data underwent quality control with fastp (v0.23.4), followed by alignment to the mouse genome (mm39) with Salmon (v1.8.0). Gene count matrices and TPM matrices were generated using tximport (v1.34.0) and TxDb.Mmusculus.UCSC.mm39.knownGene (v3.20.0) packages in R. Genes with total raw counts fewer than 10 across all samples were excluded prior to downstream analyses. Differential expression analysis was performed using the DESeq2 package (v1.46.0) in R with Wald test and Benjamini-Hochberg correction for multiple testing. Gene annotation was performed with org.Mm.eg.db package (v3.20.0) in R. We evaluated TFBS enrichment on promoter sequences defined as ±1,000 bp from the TSS, obtained from the UCSC Genome Browser (mouse genome, GRCm39), using TFBStools (v1.44.0) and JASPAR2022 (v0.99.8) packages in R. Over representation analysis (ORA) for GO terms were conducted using clusterProfiler package (v4.14.6) in R, with significant terms defined as those with adjusted *p*-value < 0.05 and *q*-value < 0.1. RRHO analysis was performed using RRHO2 package (v1.0) in R.

### Measurement of the Transcription factor activity

The endogenous activities of multiple TFs in cultured astrocytes expressing TF activity reporters were measured as previously established (28), but this study used plasmids listed above (pTFRep1–6) instead of viral particles. Briefly, TF activity was quantified as the ratio of mRNA expression levels between reporter gene and corresponding reference gene derived within a single TF activity reporter construct. This procedure controls for the variation in transfection efficiency. The reference genes are constitutively expressed under the control of a phosphoglycerate kinase (PGK) promoter, while the expression of reporter genes is induced by a minimal promoter with TF binding sites (TFBSs) for the transcription factor of interest. By varying the combinations of TFBSs and paired reference and reporter gene sets, we quantified transcription factor activity from individual samples. Total RNA was extracted using MagExtractor (Toyobo, #NPK-201F) and converted to cDNAs using ReverTra Ace qPCR RT Master Mix with gDNA Remover (Toyobo, #FSQ-301). Quantitative PCR (qPCR) was performed using TB Green Premix Ex Taq II FAST qPCR (Takara, #RR830A) on CFX384Touch (BioRad). Each reporter and reference gene expression was measured by quantitative RT-PCR using specific primer sets described in a previous study (28). Absolute quantification was performed by comparison with standard plasmids of known concentrations. TF activities were normalized to the mean activity of each TF in control samples, enabling direct comparison across hM4Di, hM3Dq, rM3Ds, and control conditions. In this study, the term “TF activity” refers to the ratio of reporter gene expression to that of the corresponding reference gene, and is presented as a relative measure across the experimental conditions. A total of 15 TF activity reporters were used, with up to 6 plasmids used per experiment.

### LASSO regression modeling

For each gene, RNA-seq–derived expression levels were quantified as log (TPM + 1). The resulting expression matrix was standardized separately for each gene. Genes exhibiting near-zero variance across samples were then filtered out, resulting in a final set of 16,639 genes that were individually subjected to model fitting. For each gene, expression levels across 12 samples (*n* = 3 biological replicates for each experimental condition) served as the response variables. To account for variability in TF activity measurements, we employed a subsampling approach. For each TF, 3 activity values were randomly sampled from the 9–12 available experimental measurements per condition. The sampled TF activities were served as predictor variables and were paired with the corresponding gene expression data from the same experimental condition. Multiple linear regression models with L1 regularization (LASSO) were then constructed using glmnet package (v4.1.10) in R, which internally standardized predictor variables separately for each TF. This procedure was repeated 200 times per gene. Model performance was evaluated by leave-one-out cross-validation (LOOCV). The regularization parameter was optimized to minimize the mean squared error during cross-validation. Genes were classified as “explained” by the model if their mean training *R*² across 200 iterations exceeded a predefined threshold.

### Astrocyte intracellular Ca^2+^ imaging and data analysis

Three days before the live imaging, cells were treated with DCZ (8 μM) or PBS (vehicle) for 90 min, after which the medium was replaced with fresh DCZ-free medium. Live imaging was performed at 37°C using inverted fluorescence microscope (Axiovert A1 MAT, Zeiss) equipped with a Colibri 7 system (Zeiss) and a stage heater (Tokai Hit). Imaging was conducted without CO_2_ supply. HEPES (40 mM, pH = 7.4; Nacalai tesque) was added to each well at least 5 min prior to recording to prevent pH change during imaging. Fluorescent signals were acquired at 10 Hz using a 10× objective lens. A droplet of ATP (final 100 μM; Fujifilm-WAKO, # 017-09673) or PBS was applied during the imaging at specified time points. Analyses of time-lapse image series were performed using ImageJ (NIH). Regions of interest (ROIs) corresponding to individual astrocytes and spatially distributed background regions were manually defined, and raw fluorescence intensity traces were extracted for each ROI. Using dplyr (v3.20.0) and tidyr (v1.3.1) packages in R, the mean background fluorescence at each time point was calculated and subtracted from all raw traces for each frame. Time traces of fluorescence intensity were converted to ΔF/F (calculated as (F − F_0_) / F_0_). F_0_ represents the baseline fluorescence, defined as the average signal during the 0–60 s period. Using slider package (v3.20.0) in R, the ΔF/F traces were smoothed with a time-window of 5. For each cell, exponential fluorescence decay associated with photobleaching was assessed using base R functions, and bleaching correction was selectively applied to cells exhibiting significant decay. After bleaching correction, calcium response quantification from ΔF/F traces was conducted. Responsive cells were defined as those with peak ΔF/F > 0.7. The integrated area-under-the-curve (AUC) was measured using pracma (v2.4.4) and dplyr (v 3.20.0) packages in R.

### Quantifications and statistical analysis

All statistical analyses were performed using R (v4.3.2 or v4.4.2). For a series of transcriptomic analyses, we used a hypergeometric distribution analysis and linear mixed-effects models. TF activity analysis was performed using one-way ANOVA followed by Tukey’s post hoc test. Pearson correlation coefficient was used in LASSO modeling. For Ca^2+^ imaging analyses, Fisher’s exact test and unpaired two-tailed *t*-test were used. Changes in astrocyte size following DREADD activation were assessed using one-way ANOVA. Numbers of *n* represent the biological replicates.

### Data and materials availability

Data and materials are available from the corresponding author upon reasonable request.

## Acknowledgments

This research was supported by JSPS/MEXT KAKENHI (JP25H010370, JP24H021460, JP24H012180 to K.A.), Tohoku University Research Program “Frontier Research in Duo” (Grant No. 2101 to K.A.), Suzuken Memorial Foundation (K.A.), Takeda Science Foundation (K.A.), and Asahi Glass Foundation (K.A.). We thank Abe lab members for kind support during this study. We also thank Dr. Ko Matsui for comments on the draft.

## Author Contributions

M.S. and K.A. designed research; M.S. and K.A. performed research; M.S. analyzed data; H.Y. contributed reagents/analytic tools; and M.S. and K.A. wrote paper.

